# A Consequence of Immature Breathing induces Persistent Changes in Hippocampal Synaptic Plasticity and Behavior: A Role of Pro-Oxidant State and NMDA Receptor Imbalance

**DOI:** 10.1101/2023.03.21.533692

**Authors:** Alejandra Arias-Cavieres, Alfredo J. Garcia

**Affiliations:** Institute for Integrative Physiology, The University of Chicago; Grossman Institute for Neuroscience, Quantitative Biology & Human Behavior, The University of Chicago; Department of Medicine, Section of Emergency Medicine, The University of Chicago

**Keywords:** apnea, prematurity, oxidative stress, hippocampus

## Abstract

Underdeveloped breathing results from premature birth and causes intermittent hypoxia during the early neonatal period. Neonatal intermittent hypoxia (nIH) is a condition linked to the increased risk of neurocognitive deficit later in life. However, the underlying mechanistic consequences nIH-induced neurophysiological changes remains poorly resolved. Here, we investigated the impact of nIH on hippocampal synaptic plasticity and NMDA receptor (NMDAr) expression in neonatal mice. Our findings indicate that nIH induces a pro-oxidant state, leading to an imbalance in NMDAr subunit composition that favors GluN2A over GluN2B expression, and subsequently impairs synaptic plasticity. These consequences persist in adulthood and coincide with deficits in spatial memory. Treatment with the antioxidant, manganese(III) tetrakis(1-methyl-4-pyridyl)porphyrin (MnTMPyP), during nIH effectively mitigated both immediate and long-term effects of nIH. However, MnTMPyP treatment post-nIH did not prevent the long-lasting changes in either synaptic plasticity or behavior. Our results underscore the central role of the pro-oxidant state in nIH-mediated neurophysiological and behavioral deficits and importance of stable oxygen homeostasis during early life. These findings suggest that targeting the pro-oxidant state during a discrete window may provide a potential avenue for mitigating long-term neurophysiological and behavioral outcomes when breathing is unstable during early postnatal life.

**Highlights:**

- Untreated immature breathing leads neonatal intermittent hypoxia (nIH).
- nIH promotes a pro-oxidant state associated with increased HIF1a activity and NOX upregulation.
- nIH-dependent pro-oxidant state leads to NMDAr remodeling of the GluN2 subunit to impair synaptic plasticity.
- Impaired synaptic plasticity and NMDAr remodeling caused by nIH persists beyond the critical period of development.
- A discrete window for antioxidant administration exists to effectively mitigate neurophysiological and behavioral consequences of nIH.

## Introduction

Human neonates are born prematurely (less than 37 weeks of gestation) are at an increased risk of Apneas of Prematurity, a condition that results in breathing instabilities causing intermittent hypoxemia and subsequently, oxidative stress (Di Fiore & Raffay, 2021; Di Fiore & Vento, 2019). While instabilities due to immature breathing resolve with continued postnatal development, the occurrence of oxidative stress during early life is hypothesized to be a principal contributor to causing disturbances in the neonatal brain (Panfoli et al., 2018) and the emergence of neurobehavioral deficits associated with intellectual disability and autism spectrum disorder in humans (Poets, 2020). Indeed, oxidative stress has been documented in animal models exposed to perinatal IH (Garcia et al., 2016; Souvannakitti, Kumar, Fox, & Prabhakar, 2009; Souvannakitti et al., 2010). Additionally, perinatal IH also causes anatomical and neurophysiological changes that are associated with deficit in affective and cognitive behaviors later in life (Cai, Tuong, & Gozal, 2011; Goussakov, Synowiec, Yarnykh, & Drobyshevsky, 2019; Vanderplow et al., 2022). However, extent to which the pro-oxidant state contributes to neurophysiological and behavioral changes caused by nIH remains poorly resolved.

During normal postnatal development, NMDAr subunit composition changes where GluN2A subunit expression progressively increases and surpasses GluN2B to become the predominant GluN2 isoform in adulthood (Paoletti, Bellone, & Zhou, 2013). As GluN2 subunit composition is an important determinant to both biophysical and downstream signaling properties of the NMDAr (Hansen et al., 2018; Paoletti et al., 2013; Paoletti & Neyton, 2007), disruption to the normal transition of GluN2 subunit composition may impact neurophysiological properties and neurocognition later in life.

Here, we characterize the immediate and long-term consequences of nIH on the hippocampus and behavioral performance related to spatial learning and memory. Here we show that the IH-dependent pro-oxidant state is linked to remodeling of GluN2 subunit composition and is associated with impaired NMDAr-dependent synaptic plasticity in the neonatal hippocampus. These nIH-dependent phenomena persist later in life and coincide with deficits related to spatial memory. While MnTMPyP treatment during IH prevents the emergence of these phenomena, antioxidant treatment after IH is ineffective. In addition to demonstrating a previously undescribed role for nIH-dependent oxidative to stress perturb the normal development trajectory of NMDAr-dependent physiology and behavior later in life, this study identifies an interventional window effective for mitigating the impact of nIH-dependent oxidative stress in promoting long-lasting consequences on neurophysiology and behavior.

## Materials and Methods

### Study Approval

All animal protocols were approved by the Institutional of Animal Care and Use Committee at the University of Chicago, in accordance with National Institute of Health guidelines.

### Animals

Mice were housed in AAALAC-approved facilities with a 12 hour/12-hour light-dark cycle and ad *libitum* to food and water. Mice used in this study were from a C57BL/6 background. To allow ad *libitum* access for nutrition, pups of both sexes and their dam were exposed together to either room air (control), nIH, nIH with daily saline administration (nIH_saline_) or nIH with daily MnTMPyP administration (nIH_Mn_). We examined age-matched control adult mice (P55 to P60; Adult_control_), nIH mice after approximately six weeks (42 ± 4 days) of recovery in room air recovery (Adult_nIH_), nIH_Mn_ mice after six weeks recovery in room air (Adult_nIH-Mn_), and nIH mice that received approximately six weeks (42 ± 4 days) of MnTMPyP after IH exposure (Adult _Rec-Mn_). Body mass immediately following IH and six weeks following IH exposure were similar to age-matched controls (**Supplement 1**). Mice treated with MnTMPyP (Enzo Life Sciences, Cat #ALX-430–070) received a single daily dose (5mg/kg i.p.) at the beginning of each day for the aforementioned treatment regimens.

### Intermittent hypoxia exposure

As previously described (Garcia et al., 2016), the nIH paradigm was executed during the light cycle and lasted for 8±1 hours per day for ten consecutive days. Exposure to nIH began on P4 to P5. A single hypoxic cycle was achieved by flowing 100% N_2_ into the chamber for approximately 60 sec. This created a hypoxic environment where the nadir O_2_ chamber reached 4.5 ± 1.5% for approximately 10 seconds immediately followed by an air break (19 ± 2.5% O_2_; 300 sec).

### Slice Preparation for Electrophysiology

Coronal hippocampal slices were prepared from mice either 24 hours following the end of nIH (P14 to P15) or approximately six weeks nIH after (P55 to P60). Mice were anesthetized with isoflurane and euthanized by rapid decapitation. The brain was rapidly harvested and blocked, rinsed with cold artificial cerebrospinal fluid (aCSF) and mounted for vibratome sectioning. The mounted brain tissue was submerged in aCSF (4°C; equilibrated with 95% O_2_, 5% CO_2_) and coronal cortico-hippocampal brain slices (350 µm thick) were prepared. Slices were immediately transferred into a holding chamber containing aCSF equilibrated with 95% O_2_, 5% CO_2_ (at 20.5±1°C). Slices were allowed to recover a minimum of one hour prior to the transfer into recording chamber and were used up to eight hours following tissue harvest. The composition of aCSF (in mM):118 NaCl, 10 Glucose, 20 sucrose, 25 NaHCO3, 3.0 KCl, 1.5 CaCl_2_, 1.0 NaH_2_PO_4_ and 1.0 MgCl_2_. The osmolarity of aCSF was 305-315 mOsm/L and equilibrated with 95% O_2_/5% CO_2_, the pH was 7.42 ± 0.2. Recordings were made at 30 ± 1*_*C in 95% O_2_ 5% CO_2_.

### Extracellular recording of the field excitatory postsynaptic potential (fEPSP)

The extracellular recording of the fEPSP was established in aCSF (31.0 ± 2°C, equilibrated with 95% O_2_ 5% CO_2_) superfused and recirculated over the preparation. The stimulation electrode, a custom constructed bipolar electrode composed of twisted Teflon coated platinum wires (wire diameter:127 µm, catalog number 778000, AM Systems.), was positioned in the Schaffer Collateral and recording electrode (<2 M*Ω*) was placed into the *stratum radiatum* of the CA1. The intensity of the electrical current (100-400µA; 0.1-0.2 ms duration) was set to the minimum intensity required to generate the 50% maximal fEPSP.

### The fEPSP was evoked every 20s

After 10 minutes of recording the baseline fEPSP, LTP was induced using Theta Burst Stimulation (TBS: four trains of 10 bursts at 5 Hz, each burst was comprised four pulses at 100 Hz; LTP_TBS_). Following stimulation, recordings continued for up to one hour. The fEPSP slope was normalized to baseline values. D-L AP5 (50 µM, Sigma-Aldrich, Cat#A5282) was used to block NMDAr; TCN-213 (5 µM, Tocris, Cat#4163) was used to block GluN2A and ifenprodil (5 µM, Tocris, Cat#0545) was used to block GluN2B. The current stimulus was set at the minimum current value (150 to 250 µA) required to evoke the initial fEPSP at 50% the maximal value. Recordings were made using either a Multiclamp 700B (Molecular Devices, San Jose, CA, USA) or using a differential amplifier (AM system, Washington, DC, USA).

### Nuclear Western Immunoblot Assay

Entire hippocampus mouse was rapidly dissected, and samples were by homogenizing using N-PER (Thermo Fisher Scientific, Cat#87792) in cold ice by following manufacturer instructions. Briefly, cytoplasmic fragment was obtained by homogenizing tissue using a tissue grinder and then by pipetting in cytoplasmic extraction buffers. After isolation of cytoplasmic fragment, the insoluble pellet that contains nuclear proteins was suspended in nuclear extraction buffer and separated by centrifugation. Halt Protease Inhibitor (Thermo Fisher Scientific, Cat#78429) was added into cytoplasmic and nuclear extraction buffers to prevent protein degradation. Samples were boiled for 15 min in loading buffer (Bio-Rad, Hercules, CA, USA) and denaturated by *β*-mercaptoethanol at 75°C before loading 50-60 µg protein onto 4-20% Mini-PROTEAN TGX Stain-FreeTM Protein Gels (Bio-Rad, Hercules, CA, USA) and electrophoresed at 120 V for 120 min, then gels were transferred to PVDF membrane (Bio-Rad) using Transfer-Blot Turbo System (BIO-RAD, Hercules, CA, USA). Later, membranes were subsequently blocked for 2 h at room temperature with 5% bovine serum albumin (BSA) (Sigma-Aldrich, MN, USA) in Tris buffered saline (TBS) (Bio-Rad, Hercules, CA, USA). Membranes were incubated under constant shaking with primary antibodies: monoclonal mouse anti-HIF1a (1:500; Abcam Cat# ab1, RRID:AB_296474) and monoclonal rabbit anti-TBP (1:2000; Cell Signaling Technology Cat# 44059, RRID:AB_2799258). After washing three times with TBS-Tween 0.3% for 20 min, the membranes were incubated for 1.5 hr at room temperature with appropriate secondary antibodies. Finally, the membranes were washed three times with TBS-Tween 0.3% for 20 min and immunoreactive proteins were detected with Super Signal^TM^ West Femto reagents according to the manufacturer instructions (Thermo Fisher Scientific, Cat#34095). Signals were captured with the ChemiDoc system (Bio-Rad, Hercules, CA, USA). The IMAGE J image program (National Institutes of Health, USA) was used to quantify optical band intensity.

### Whole cell Western Immunoblot Assay

Entire hippocampus mouse was rapidly dissected, and samples were by homogenizing using M-PER^TM^ (Thermo Fisher Scientific, Cat#78501) Halt Protease Inhibitor (Thermo Fisher Scientific, Cat#78429) in cold ice. Samples were centrifuged at 12 rpm for 15 min at 4°C and the pellet was discarded. Samples were boiled for 15 min in loading buffer (Bio-Rad, Hercules, CA, USA) and denaturated by *β*-mercaptoethanol at 75°C before loading 20-30 µg protein onto 4-20% Mini-PROTEAN TGX Stain-FreeTM Protein Gels (Bio-Rad, Hercules, CA, USA) and electrophoresed at 120 V for 120 min, then gels were transferred to PVDF membrane (Bio-Rad) using Transfer-Blot Turbo System (BIO-RAD, Hercules, CA, USA). Later, membranes were subsequently blocked for 2 h at room temperature with 5% non-fat milk (Bio-Rad, Hercules, CA, USA) or 5% bovine serum albumin (BSA) (Sigma-Aldrich, MN, USA) in Tris buffered saline (TBS) (Bio-Rad, Hercules, CA, USA). Membranes were incubated under constant shaking with primary antibodies: anti rabbit GluN1 (1:2000; Abcam Cat# ab109182, RRID:AB_10862307) anti-rabbit GluN2A (1:2000; Cell Signaling Technology Cat# 4205, RRID:AB_2112295), anti-rabbit GluN2B (1:2000; Cell Signaling Technology Cat# 14544, RRID:AB_2798506), anti-rabbit NOX2 (1:2000; Abcam Cat# ab129068, RRID:AB_11144496) anti-rabbit NOX4 (1:500; Novus Cat# NB110-58851B, RRID:AB_1217375 and anti-mouse GAPDH (1: 10.000; Abcam Cat# ab8245, RRID:AB_2107448). Incubations were performed at 4*_*C overnight in 5% non-fat milk or BSA.

After washing three times with TBS-Tween 0.2% for 15min, the membranes were incubated for 1.5 h at room temperature with appropriate secondary antibodies. Finally, the membranes were washed three times with TBS-Tween 0.2 % for 15 min and immunoreactive proteins were detected with enhanced chemiluminescence reagents according to the manufacturer instructions (Bio-Rad, Hercules, CA USA). Signals were captured with the ChemiDoc system (Bio-Rad, Hercules, CA, USA). The IMAGE J image program (National Institutes of Health, USA) was used to quantify optical band intensity.

### TBARS assay

Whole cell protein lysates were isolated from entire hippocampal tissues using RIPA buffer (Thermo Fisher Scientific, Catalogue No. R0278) in the presence of protease and phosphatase inhibitors (Thermo Fisher Scientific, Catalogue No. 78429) in cold ice. Protein lysates were immediately processed and stored at −80°C until used. The amount of lipid-peroxidation was determined using a TBARS Assay Kit (Cayman Chemical, Cat#10009055), per manufacturer instructions. Absorbance was measured between 530-540 nm using a plate reader. The analysis for determine MDA values was made according with per manufacturer instructions.

### Barnes maze

The Barnes maze performed in using a custom made opaque white circular acrylic platform (92.4 cm in diameter) with 20 equidistant holes (5.08 cm in diameter and 2.54 cm from the edge). The platform was elevated (30 cm from the floor) ground and surrounded by four identical walls (27.94 cm high). By default, each hole was closed with a fixed piece of opaque acrylic that could be removed to lead to a dark exit box. Lighting was achieved through diffuse overhead fluorescent lighting such that all holes were equally lit. An overhead camera was suspended above the maze. Data collection and posthoc analysis was performed using CinePlex Video Tracking System (Plexon, Dallas, TX).

As previously described (Arias-Cavieres et al., 2020), the task was performed using a four-day protocol consisting of one training trial per day for three consecutive days and a probe trial on the fourth day. Barnes Maze began after 6 weeks exposure to IH with respective controls run at the same time. For the training trials, all, but one of the holes (exit hole), were closed. An exit box with a small ramp was placed directly underneath the exit hole. Animals were given a maximum of six minutes to locate the exit and if unable to locate the exit, they were gently guided to the exit. If the mouse found and entered the exit before the six minutes were over, the trial ended at the time that the mouse entered the exit, and the mouse was promptly returned to its home cage. During the probe trial, all holes were closed, and the animal was given six minutes to explore the maze. The entire arena was sanitized in-between trials.

Entry probability for each hole during the probe trail was calculated by the following:

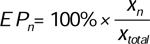

where *EP_n_* is the entry probability for hole n; *x_n_* = number of entries into hole n; and *x_total_* is sum of entries across all holes during the probe trial.

### Object Location Task

The Object Location task was performed using a modified protocol described by (Wimmer, Hernandez, Blackwell, & Abel, 2012). The protocol was modified for mice. The procedure included three phases: open field; familiarization, and object location. Each phase was performed in acrylic open arena (W: 33 cm, L:36 cm, H: 33 cm). During the open field phase, the mouse was habituated and allowed to freely explore the arena for 10 min. The familiarization consisted of three consecutives (session duration: 5 min; intersession duration: 5 min), where the mouse was placed in the arena and allowed to explore two different objects (i.e., object A and object B). The time exploring both objects were recorded. The object location phase occurred 24 hr following familiarization where one object presented in the familiarization was repositioned (B). The times exploring the relocated object (object B) was used to compared with the other group. To eliminate odor cues between trials, the experimental apparatus and all objects were cleaned with 75% ethanol after each trial. Mice behavior was recorded with a video camera positioned over the behavioral apparatus and the collected videos were analyzed with the ANY-MAZE software (Stoelting Co., Wood Dale, IL, USA).

### Statistical Analysis

Statistics were performed using Prism 6 (GraphPad Software, Inc.; RRID: SCR_015807) and results were plotted in either Prism 6 or Origin (Origin Labs;2018b). Comparisons between two groups were conducted using unpaired two-tailed *t*-test with Welch’s correction or paired comparisons, where appropriate. The equality of variances between two groups was determined with an F test. Analyses involved comparisons beyond more than two groups were assessed by a one-way ANOVA of means followed by a *posthoc* Bonferroni’s multi-comparison test. Unless otherwise stated, data are presented as mean ± S.E.M, and where appropriate, individual responses overlaid over box plots. The upper and lower limit of each box represented 25^th^ and 75^th^ percentile of the cohort, respectively, while the inter-limit line within the box represented median. The Box plot error bars represented maximum and minimum values in the data set. Significance was defined as P<0.05.

## Results

### nIH promotes a pro-oxidant state, increases nuclear HIF1a upregulates NADPH Oxidase and promotes a pro-oxidant state in the neonatal hippocampus

Measurements of malondialdehyde (MDA) content in neonatal hippocampal homogenates revealed that MDA levels were increased with nIH (Figure 1A, control: 7.2 ± 0.82 nmol • mg^-1^ protein and nIH:18.59 ± 1.65 nmol • mg^-1^ protein, P=0.004; N=6 preparations per condition). This was accompanied by greater nuclear content of the pro-oxidant transcription factor, HIF1a (Figure 1B, control= 1.06 ± 0.04 and nIH=1.59±0.12, P=0.010; N=5 per condition) and increased expression of two isoforms of the pro-oxidant enzyme NADPH oxidase (NOX), NOX-2 (Figure 1C control= 1.02±0.02 and nIH=1.31±0.12; P=0.045; N=6 per condition) and NOX-4 (Figure 1D, NOX-4: control= 0.96±0.03 and nIH=1.30±0.07; P=0.003; N=5 per condition).

**Figure 1:**
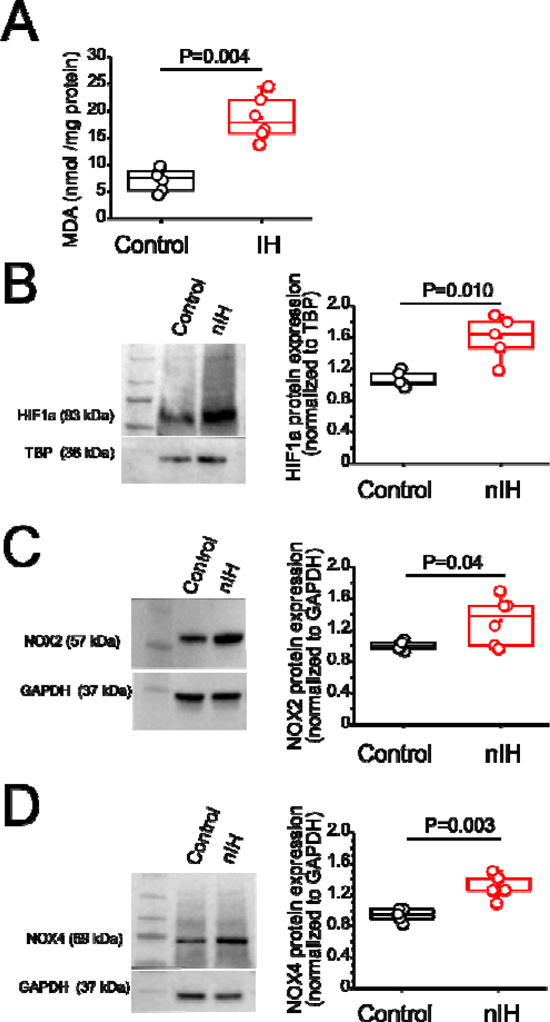
nIH promotes a pro-oxidant state that coincides with increased nuclear HIF1a and NOX4 isoforms in the neonatal hippocampus. **A**. Malondialdehyde (MDA) content was measured in hippocampal homogenates from control and exposed to nIH (two tailed t-test, t=6.16; df=7.33; P=0.004). **B**. (*left*) Representative blot of nuclear HIF1a from mice unexposed and exposed to nIH. (right). Quantification of nuclear HIF1a expression from neonate control and IH exposed mice (two tailed t-test, t=3.98; df=4.98; P=0.010). **C**. (*left*) Immunoblot of NOX2. (*right*) Significant differences were found in hippocampal homogenate from nIH vs control. (two tailed t-test, t=2.59; df=5.3; P=0.04). **C**. (left) Representative image of NOX4. (*right*) Comparison of NOX4 expression between control and mice exposed to nIH (two tailed t-test, t=4.53; df=6.01; P=0.003). **D**. The analysis was performed for A, B, C and D using unpaired two-tailed t-test with Welch’s correction.

### Attenuation of synaptic plasticity by nIH is associated with changed NMDAr subunit composition

In the adult rodent IH-dependent increased nuclear HIF1a and the shift toward a pro-oxidant state is associated with impaired NMDAr-dependent synaptic plasticity (Arias-Cavieres, Fonteh, Castro-Rivera, & Garcia, 2021; Arias-Cavieres et al., 2020). Therefore, we next characterized how nIH impacted LTP evoked by theta-burst stimulation (LTP_TBS_) in neonatal hippocampal slices from control and nIH mice. LTP_TBS_ was evoked in area CA1 in both control (Figure 2A, black, n=7 slices; N=6 mice) and nIH (Figure 2A, red, n=6; N=5) and blockade of NMDAr, using AP5 [50 µM], prevented LTP_TBS_ in both groups (Figure 2A, control: green, n=4, N=4; nIH: blue, n=4, N=4). However, the magnitude of LTP_TBS_ was smaller in nIH slices as compared to control (Figure 2A, control: 77.02±5.9% versus nIH: 35.10±6.8% over baseline, P=0.009). Such a difference may have resulted from an overall downregulation in NMDAr expression or a change in receptor subunit composition.

**Figure 2:**
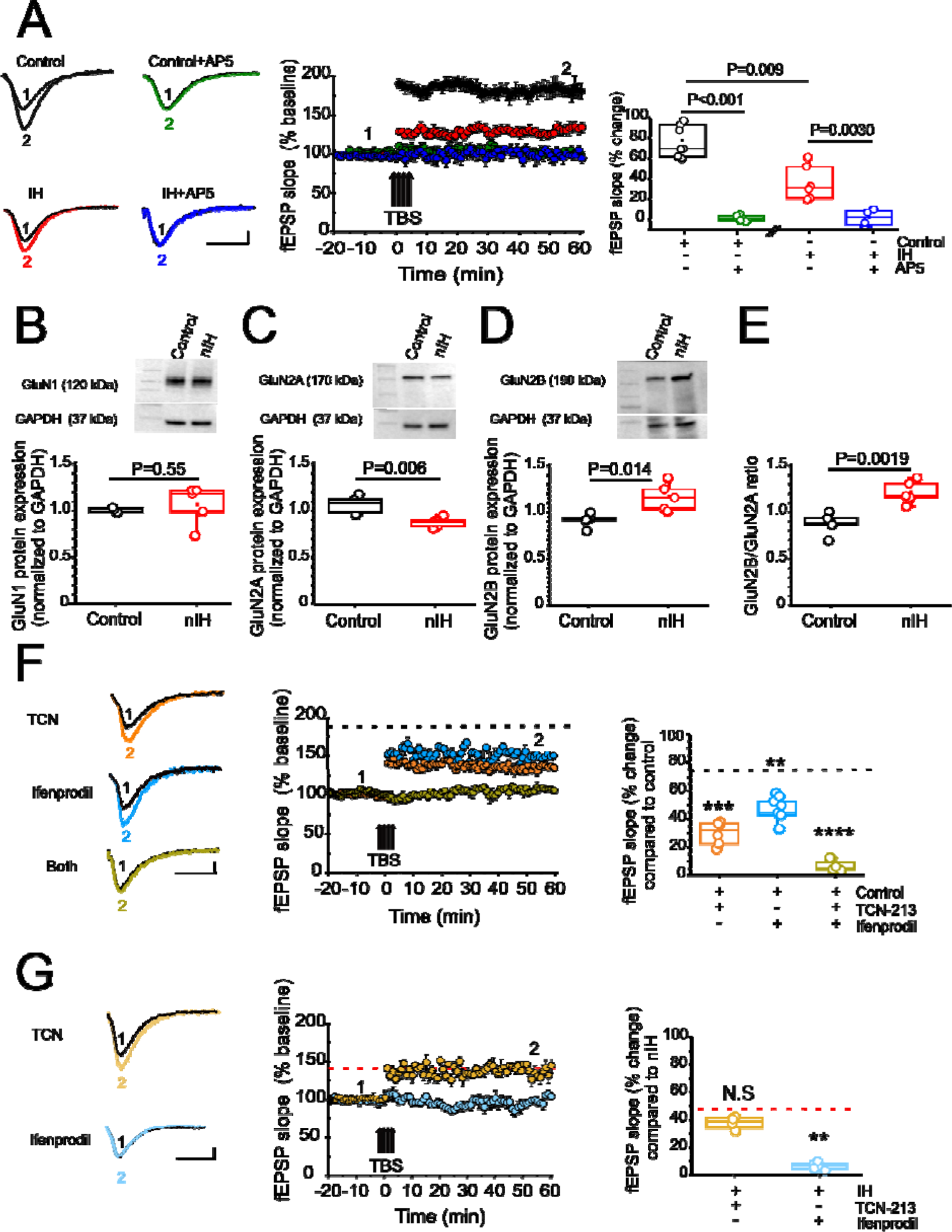
nIH suppresses LTP and changes the sensitivity of LTP to TCN-213 and ifenprodil. **A**. (*left*) Representative traces of the evoked fEPSP from control (black), control+AP5 [50µM] (green), nIH (red) and nIH +AP5 [50µM] (blue) in baseline conditions prior to TBS (1) and following TBS (2). (*middle*) Mean fEPSP slope plotted as a function of time relative to the slope before TBS in control, control+AP5, nIH and nIH+AP5. (*right*) fEPSP slope represented as percent change from baseline at 60 min after TBS in control slices vs control+AP5 [50 µM]. **B**. (*top*) Representative image for GluN1. (bottom) The graph shows GluN1 did not change protein expression after nIH exposure compared with control. (two tailed t-test, t=0.63; df=4.1; P=0.55). **C**. (top) Western blot picture for GluN2A. (bottom) Comparison for both conditions show GluN2A decreased the protein content levels after nIH. (two tailed t-test, t=4.017; df=6.43; P=0.006). **D**. (*top*) Representative immunoblot image for GluN2B. (bottom) Comparison of both conditions show increased GluN2B levels after nIH (two tailed t-test, t=3.43; df=5.78; P=0.014). **E**. GluN2B/GluN2A comparison ratio. **F**. (*left*) Representative traces of the evoked fEPSP control +TCN [5 µM] (orange), control + ifenprodil [5 µM] (cyan) and control + both drugs (olive) in baseline conditions prior to TBS (1) and following TBS (2). (*middle*) Mean fEPSP slope plotted as a function of time relative to the slope before TBS and (*right*) Comparison of fEPSP slope represented as percent change from baseline at 60 min after TBS (one way ANOVA, F_(3,20)_=43.52; P<0.001). Black dashed line represents the mean slope of the fEPSP 60 min following TBS in control slices from Figure 2A. **G**. (*left*) Representative traces of the evoked fEPSP from nIH+TCN [5 µM] (dark yellow) and nIH +ifenprodil [5 µM] (light blue) in baseline conditions prior to TBS (1) and following TBS (2). (*middle*) Mean fEPSP slope plotted as a function of time and relative to slope before TBS (*right*) fEPSP slope represented as percent change from baseline at 60 min after TBS (two tailed t-test, t=12.46; df=4.59; P=0.001). Red dashed line represents the mean slope of the fEPSP 60 min following TBS in nIH slices from Figure 2A. Scale bars for A, F and G: 10 msec x 0.2 mV. The box-plot parameters indicate mean ± S.E. The analysis was performed for A to E using unpaired two-tailed t-test with Welch’s correction and for F and G the analysis was performed using one-way ANOVA followed by Bonferroni post hoc. **P<0.01, ***P<0.001 and ****P<0.0001).

While expression of GluN1, the obligatory subunit of the NMDAr, was similar between control and nIH (Figure 2B, control: 1.06±0.01, N=5; nIH: 1.11±0.09, N=5; P=0.55), GluN2 subunit composition appeared to be different between the groups. GluN2A expression was reduced following nIH (Figure 2C, control: 1.07 ± 0.04, N=5; nIH: 0.88 ± 0.02, N=5; P=0.036); whereas, GluN2B expression following nIH was greater that control (Figure 2D, control: 1.06±0.03; N=5; nIH: 1.38±0.09; N=5, P=0.014). These differences coincided with a larger GluN2B:GluN2A ratio following IH (Figure 2E, P=0.0019). As GluN2 subunit composition is a significant factor dictating NMDAr-dependent physiology (A. Kumar, Thinschmidt, & Foster, 2019; S. S. Kumar & Huguenard, 2003; Paoletti et al., 2013), we sought determine whether changed LTP_TBS_ following nIH was related to the changes in GluN2 subunit composition. In control slices, the magnitude of LTP_TBS_ was reduced when blocking either the GluN2A containing NMDAr with TCN-213 (Figure 2F, orange, 29.87 ± 3.07% over baseline; n=7 slices, N=6) or the GluN2B containing NMDAr with ifenprodil (Figure 2F, cyan, 46.67 ± 3.87% over baseline; n=6 slices, N=6). When both agents were applied LTP_TBS_ was not effectively evoked (Figure 2F, olive, 6.37 ± 2.03% over baseline; n=4 slices, N=4).

Following nIH, LTP_TBS_ appeared to be unaffected by TCN-213 (Figure 2G; light yellow, 37.58 ± 2.24% over baseline; n=4 slices, N= 4 mice) when compared to treated nIH slices (from Figure 2A), yet ifenprodil appeared to prevent LTP_TBS_ following nIH (Figure 2G; light blue, 6.13 ± 1.16% over baseline, n=5 slices, N= 5). Collectively, these results suggest that NMDAr-dependent plasticity appears to be almost exclusively driven by GluN2B containing receptors and is associated with the nIH-mediated remodeling of GluN2 subunit composition to favor GluN2B expression over GluN2A.

### MnTMPyP administration during IH mitigates IH-dependent molecular, biochemical and neurophysiological changes in the neonatal hippocampus

To determine the contribution of the nIH-dependent pro-oxidant state to the observed changes in the neonatal hippocampus, we administered, the superoxide anion scavenger, MnTMPyP to a cohort of mice during nIH. While MDA content in neonatal hippocampi from the control and the cohort receiving MnTMPyP during nIH (nIH_Mn_) were similar, MDA content in nIH cohort treated with saline vehicle (nIH_saline_) was greater than control (Figure 3A; control: 9.31 ± 0.98 nmol per mg protein; nIH_saline_: 16.75 ± 1.78 nmol per mg protein; nIH_Mn_: 11.33 ± 0.93 nmol per mg protein; P=0.0011; n=5 per group). Nuclear HIF1a content (Figure 3B, control: 0.99 ± 0.07; nIH_saline_: 1.32±0.10 and nIH_Mn_: 1.03 ± 0.05, P=0.017; n=5 per group), NOX2 (Figure 3C, control: 1.03±0.03, nIH_saline_: 1.32 ± 0.051 and nIH_Mn_: 0.99 ± 0.046, P=0.024; N=6 per group) and NOX4 were increased in nIH_saline_ (Figure 3D, control: 1.02± 0.03; nIH_saline_: 1.35±0.08 and nIH_Mn_: 1.10 ± 0.01, P=0.0034; n=4 per group) and appeared to be unchanged in nIH_Mn_. GluN2A expression was also reduced in nIH_saline_ while expression appeared to be unchanged in nIH_Mn_ (Figure 3E, control: 1,08 ± 0.07; nIH_Saline_: 0.68 ± 0.06; nIH_Mn_: 0.92 ± 0.06; P=0.0086; N=6 per group).

**Figure 3:**
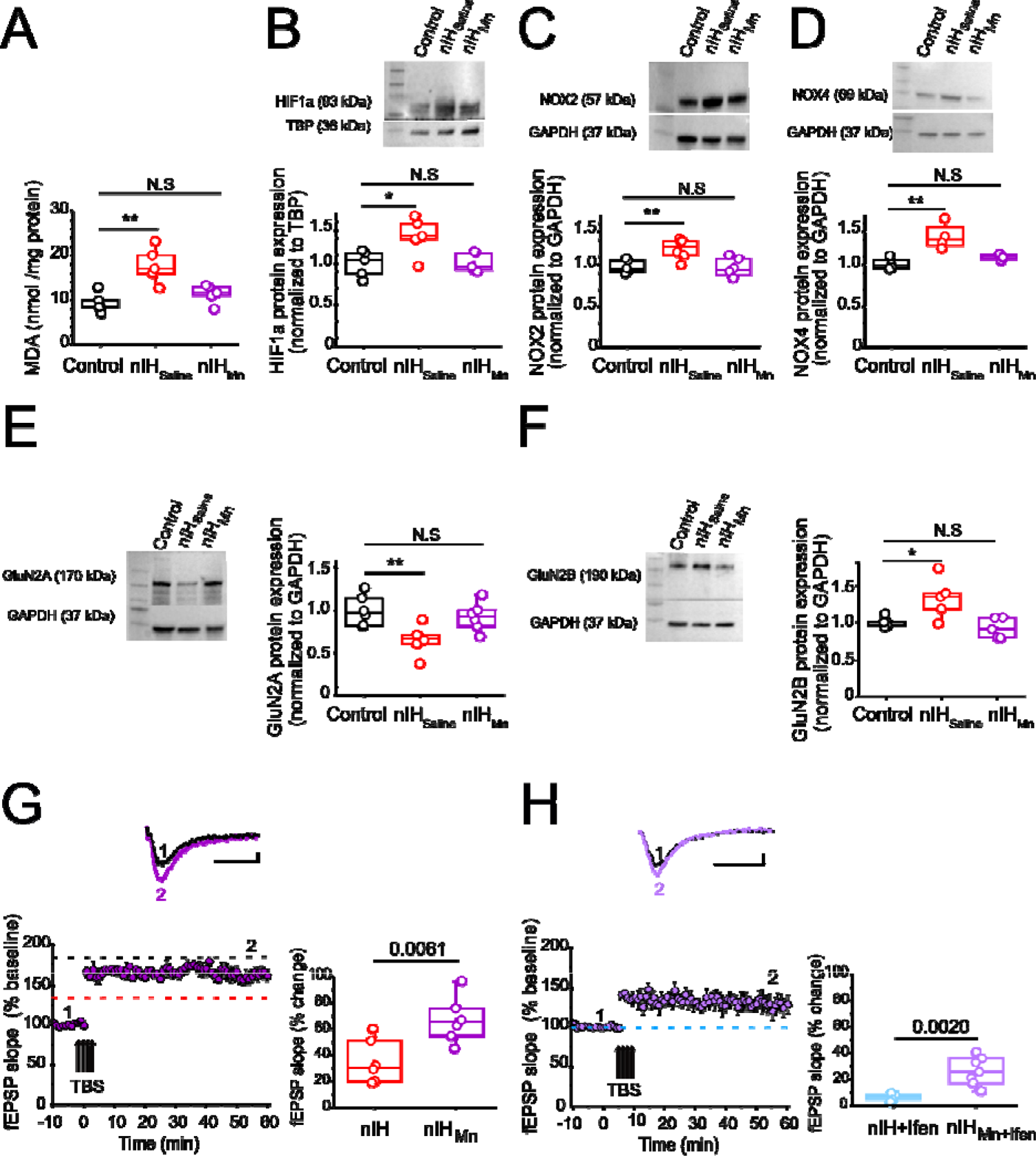
MnTMPyP administration mitigates nIH-mediated oxidative stress, increased nuclear HIF1a, NOX isoform upregulation, and impairments to LTP. **A**. Malondialdehyde (MDA) content was measured in hippocampal homogenates from control, nIH_Saline_ and 10-Mn. (one way ANOVA, F_(2,12)_=11.53; P=0.0016). **B**. (top) Representative blot of nuclear HIF1a performed from control, nIH_Saline_ and IH_10-Mn_ mice. (bottom) Quantification of HIF1a expression (one way ANOVA, F_(2,12)_=5.76; P=0.017). **C**. (top) Immunoblot of NOX2. (bottom) Significant differences was found in hippocampal homogenate from nIH_Saline_ compared to control and nIH_Mn_ (one way ANOVA, F_(2,15)_=9.26; P=0.0024). **D**. (top) Representative image of NOX4. (*bottom*) Comparison of NOX4 expression between control, nIH_Saline_ and nIH_Mn_. (one way ANOVA, F_(2,9)_=11.46; P=0.0034). **E**. (*left*) Representative blot of GluN2A from control, nIH_Saline_ and nIH_Mn_ mice. (*right*). Quantification of GluN2A expression. (one way ANOVA, F_(2,15)_=6.63; P=0.0086). **F**. (*left*) Immunoblot of GluN2B. (*right*) Significant differences were found in hippocampal homogenate from nIH_Saline_ vs control and nIH_Mn_. (one way ANOVA, F_(2,12)_=6.88; P=0.01). **G**. (top) Representative traces of evoked fEPSP from nIH_Mn_ (purple) in baseline conditions prior to TBS (1) and following TBS (2). (*left bottom*) Mean fEPSP slope plotted as a function of time relative to the slope before TBS in nIH_Mn_. (*right*) fEPSP slope represented as percent change from baseline at 60 min after TBS in nIH vs nIH_Mn_ slices. (two tailed t-test, t=3.41; df=10.61; P=0.0061). Dashed lines represent the mean slope of the fEPSP 60 min following TBS in control (black dashed line) and nIH (red dashed line) slices from Figure 2A. **H**. (top) Representative traces of the evoked fEPSP from nIH_Mn_ in presence the ifenprodil [5 µM] (light purple) in baseline conditions prior to TBS (1) and following TBS (2). (*left bottom*) Mean fEPSP slope plotted as a function of time relative to the slope before TBS in nIH_Mn_ in presence the ifenprodil. (*right*) fEPSP slope represented as percent change from baseline at 60 min after TBS in IH+ifenprodil vs nIH_Mn_ in presence of ifenprodil. (two tailed t-test, t=3.99; df=6.99; P=0.019). Blue dashed line represents the mean slope of the fEPSP 60 min following TBS in ifenprodil treated nIH slices from Figure 2G. For G and H, Scale bars 10 msec x 0.2 mV. The analysis was performed for A to F using one-way ANOVA followed by Bonferroni post hoc and for G and F, the analysis was performed using unpaired two-tailed t-test with Welch’s correction. *P<0.05, **P<0.01, and ***P<0.001.

Furthermore, while GluN2B expression was increased in nIH_saline_, GluN2B expression appeared to be unchanged in nIH_Mn_ (Figure 3F, control: 1.03 ± 0.030; nIH_saline_: 1.38± 0.12; nIH_Mn_: 1.02 ± 0.059, P=0.010; n=5 per group). Similarly, the magnitude of NMDAr-dependent in nIH_Mn_ was greater than nIH (Figure 3G, nIH_Mn_: 66.65±6.20 % over the baseline; n=7, N=5) and the sensitivity to of LTP to ifenprodil was evident when compared to nIH (Figure 3H, 33.26 ± 5.03 % over the baseline; n=7, N=5).

### IH causes persistent deficits in NMDAr-dependent synaptic plasticity and NR2 subunit remodeling that can be mitigated by MnTMyPyP administration

To assess the long-term consequences of nIH, we characterized LTP in adult mice allowed to recover from nIH for six weeks in room air (Adult_nIH_). In hippocampal slices from Adult_nIH_, the magnitude of LTP was smaller compared to that in Adult_control_ preparations (Figure 4A; Adult_control_ (black): 67.58 ± 4.08% over baseline; n=6 slices, N=6 mice; Adult_nIH_ (red): 39.47±2.76% over baseline; n=7 slices, N= 7 mice; P=0.003). To determine the physiological consequences of GluN2 levels, we explore GluN2A and GluN2B subunit contribution on LTP. To determine whether the difference in LTP magnitude observed in Adult_nIH_ hippocampal slices was related to GluN2 subunit composition, the sensitivity of LTP to TCN-213 and ifenprodil was assessed. In Adult_control_ slices, NMDAr-dependent LTP was appeared largely dependent on the GluN2A containing NMDAr and independent of GluN2B containing NMDAr as TCN-213 suppressed LTP (Figure 4B, orange, 19.53 ± 1.97% over baseline; n=4 slices, N=4 mice) while ifenprodil minimally affected LTP (Figure 4B, blue, 50.04 ± 3.59% over baseline; n=4 slices, N=4 mice). In contrast LTP from Adult_nIH_ slices appeared dependent on GluN2B containing NMDAr and independent of GluN2A containing NMDAr as ifenprodil blocked LTP (Figure 4C, blue: 6.94±1.04 over baseline; n=6, N=5) yet was insensitive to TCN-213 (Figure 4C, orange, 32.08±5.1 over baseline; n=5, N=3).

**Figure 4:**
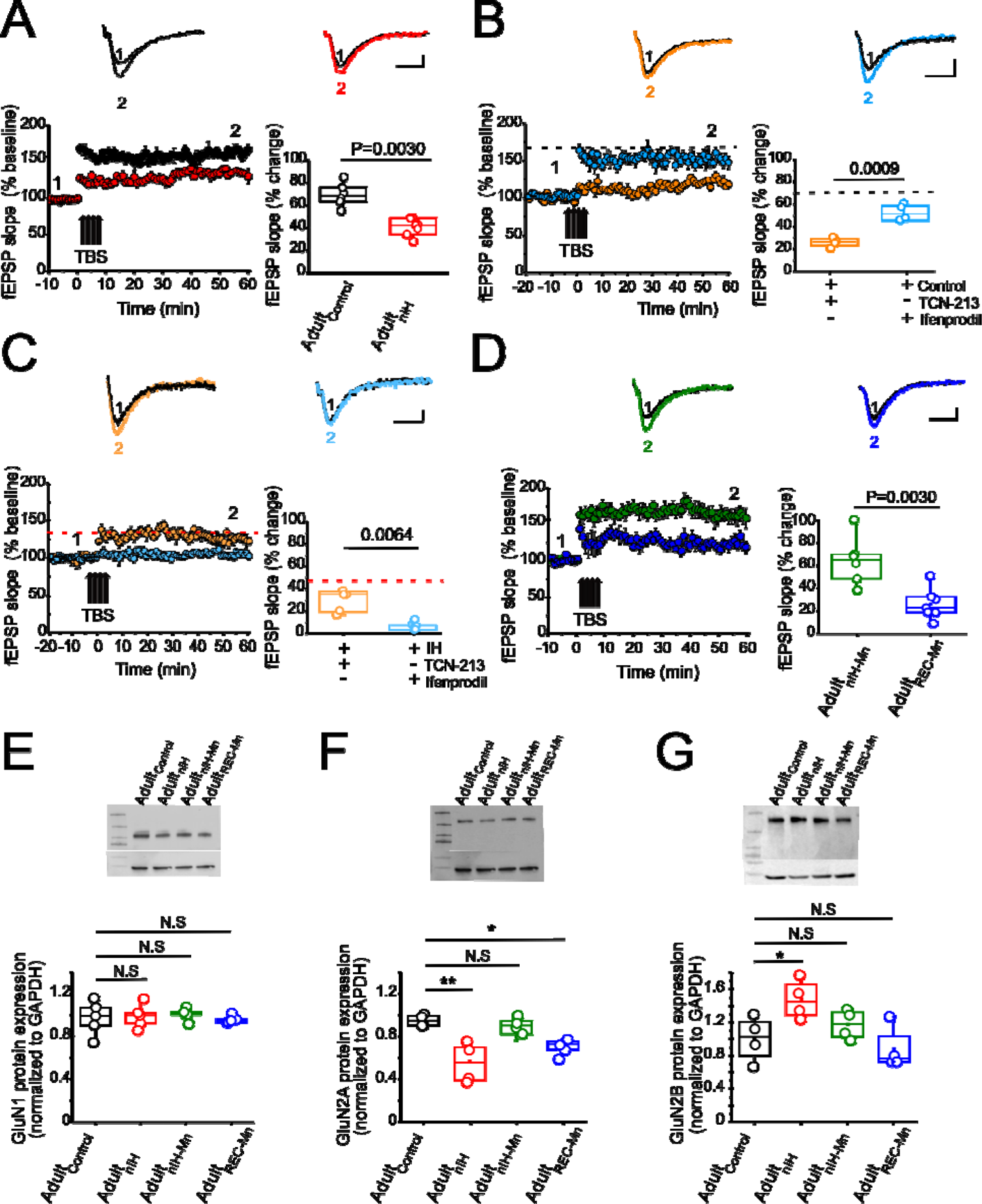
Changes in synaptic plasticity and NMDAr subunit remodeling is present in the adult hippocampus from mice exposed to nIH and can be mitigated by MnTMPyP administration during nIH. **A**. (top) Representative traces of the evoked fEPSP from adult control (black) and adult mice were exposed to neonatal IH (red) in baseline conditions prior to TBS (1) and following TBS (2). (*left bottom*) Mean fEPSP slope plotted as a function of time and relative to baseline prior to TBS. (*right bottom*) fEPSP slope represented as percent change from baseline at 60 min following TBS in adult control vs Adult_nIH_ (two tailed t-test, t=5.70; df=9.04; P=0.003). **B**. (top) Representative trace of the evoked from control+ TCN-213 [5 µM] and control + ifenprodil [5 µM] in baseline conditions prior to TBS (1) and following TBS (2). (*left bottom*) Mean fEPSP slope plotted as a function of time and relative to baseline prior to TBS. (*right bottom*) fEPSP comparison at 60 min following TBS in control +TCN and control+ ifenprodil (two tailed t-test, t=7.43; df=4.65; P=0.0009). Black dashed line represents the mean slope of the fEPSP 60 min following TBS in control slices from Figure 4A. **C**. (top) Representative trace of the evoked response from Adult_nIH_ + TCN-213 [5 µM] and Adult_nIH_ + ifenprodil [5 µM] in baseline conditions prior to TBS (1) and following TBS (2). (*left bottom*) Mean fEPSP slope plotted as a function of time relative to baseline prior to TBS. (*right bottom*) fEPSP slope represented as percent change from baseline at 60 min following TBS (two tailed t-test, t=4.72; df=4.06; P=0.0064). Red dashed line represents the mean slope of the fEPSP after 60 min following TBS in control slices from Figure 4A. **D**. (top) Representative traces of the evoked fEPSP from adult mice was exposure to neonatal IH+10 days of MnTMPyP (Adult_nIH-Mn_) and adult mice receive MnTMPyP after exposure to neonatal IH (Adult_REC-Mn_) in baseline conditions before TBS (1) and after TBS (2). (*left bottom*) Mean fEPSP slope plotted as a function of time and relative to slope before TBS in Adult_nIH-Mn_ vs Adult_REC-Mn_ (*right bottom*) fEPSP slope represented as percent change from baseline at 60 min after TBS Adult_nIH-Mn_ vs Adult_REC-Mn_ (two tailed t-test, t=3.79; df=8.16; P=0.0051). E. Representative image of GluN1. (*bottom*) Quantification shows GluN1 protein expression is not changed in Adult_nIH_, Adult_nIH-Mn_ or 10+Mn exposures compared to control (one way ANOVA, F_(3,16)=1.14_; P=0.93; N=5). **F**. (top) Representative blot of GluN2A performed from adult mice unexposed, Adult_nIH_, Adult_nIH-Mn_ or Adult_REC-Mn_. (*bottom*). Quantification of GluN2A expression from adult control, Adult_nIH_, Adult_nIH-Mn_ or 10+Mn. (one way ANOVA, F_(3,12)=8.31_; P=0.0029, N=4). **G**. (top) Immunoblot of GluN2B. (*bottom*) Significant differences were found in hippocampal homogenates from adult control, Adult_nIH_, Adult_nIH-Mn_ or Adult_REC-Mn_ (one way ANOVA, F_(3,12)=4.91_; P=0.018, N=4). The box plot parameters indicate mean ± S.E. The analysis was performed for A-D using unpaired two-tailed t-test with Welch’s correction. The analysis was performed for E-G using one-way ANOVA followed by Bonferroni post hoc. *P=0.05, **P=0.01, ***P=0.001 and N.S= no significant. Scale bars for A,B, F and G= 10 msec x 0.2 mV

As MnTMPyP prevented the immediate effects of nIH on LTP, we assessed how nIH_Mn_ and MnTMPyP administration following nIH (Adult_REC-Mn_) impacted LTP in adult hippocampal slices. The magnitude of LTP from adult mice from the Adult_REC-Mn_ was smaller in magnitude of LTP from Adult_nIH-Mn_ (Figure 4D, Adult_nIH-Mn_: green, 61.35 ±8.39% over baseline; n=6 slices, N=6 mice; Adult_REC-Mn_: blue, 24.50 ± 4.87% over baseline; n=7 slices, N=7 mice; P=0.0030).

We next examined biochemical and protein expression in the adult hippocampus. Neither HIF1a nor NOX isoforms were different from Adult_control_ in Adult_nIH_, Adult_nIH-Mn_, or Adult_REC-Mn_ (**Supplement 2**). Similarly, no differences in GluN1 subunit expression was found across experimental groups (Figure 4E, control=1.00±0.07; Adult_nIH_=0.99±0.07; Adult_nIH-Mn_=1±0.05 and Adult_REC-Mn_=0.95±0.03, N=5; P=0.93). However, differences in both GluN2A and GluN2B subunit expression were evident. In Adult_nIH_ and Adult_REC-Mn_, GluN2A expression was suppressed when compared to Adult_control_, yet no difference was evident in Adult_nIH-Mn_ (Figure 4F, Adult_control_: 1±0.05; Adult_nIH_: 0.51±0.72; Adult_nIH-Mn_: 0.92 ±0.07 and Adult_REC-Mn_: 0.80 ±0.07, N=5; P=0.0014). While GluN2B expression was increased in Adult_nIH_, no differences were observed in Adult_nIH-Mn_ and Adult_REC-Mn_ (Figure 4E, Adult_control_: 1.02 ±0.03; Adult_nIH_: 1.47±0.11, Adult_nIH-Mn_: 1.14 ±0.08, and Adult_REC-Mn_: 0.98 ±0.13, N=4; P=0.018).

### Spatial memory deficits occurring in adult mice previously exposed to nIH can be mitigated by MnTMPyP

To determine whether the persistent changes in synaptic plasticity and NMDAr subunit expression due to neonatal IH corresponded with behavioral deficits related to spatial learning and memory, we next examined the performance of control adult mice (Adult_control_) and Adult_nIH_ and in the Barnes maze and Object Location task.

In the Barnes maze, locomotor activity between control (N=14) and Adult_nIH_ (N=20) did not appear to be different as velocity and distance travelled were similar (**Supplement 3**). Moreover, training control and Adult_nIH_, exhibited progressive decreases in total latency to exit across of three training session (**Supplement 4**). During the probe trial, the initial distance (Figure 5A (*middle*); control: 0.13 ± 0.017 m versus Adult_nIH_: 0.33± 0.05 m, P=0.0019) and latency to initial entry (Figure 5A (*right*); control: 43.04± 10.44 s versus Adult_nIH_: 83.13±15.99 s, P=0.04) into the exit zone were greater in Adult_nIH_. These differences were accompanied by a smaller entry probability into the exit zone in Adult_nIH_ (Figure 5B (*right*); control: 12.63±1.42% versus Adult_nIH_: 7.14 ± 1.21%, P=0.0067).

**Figure 5:**
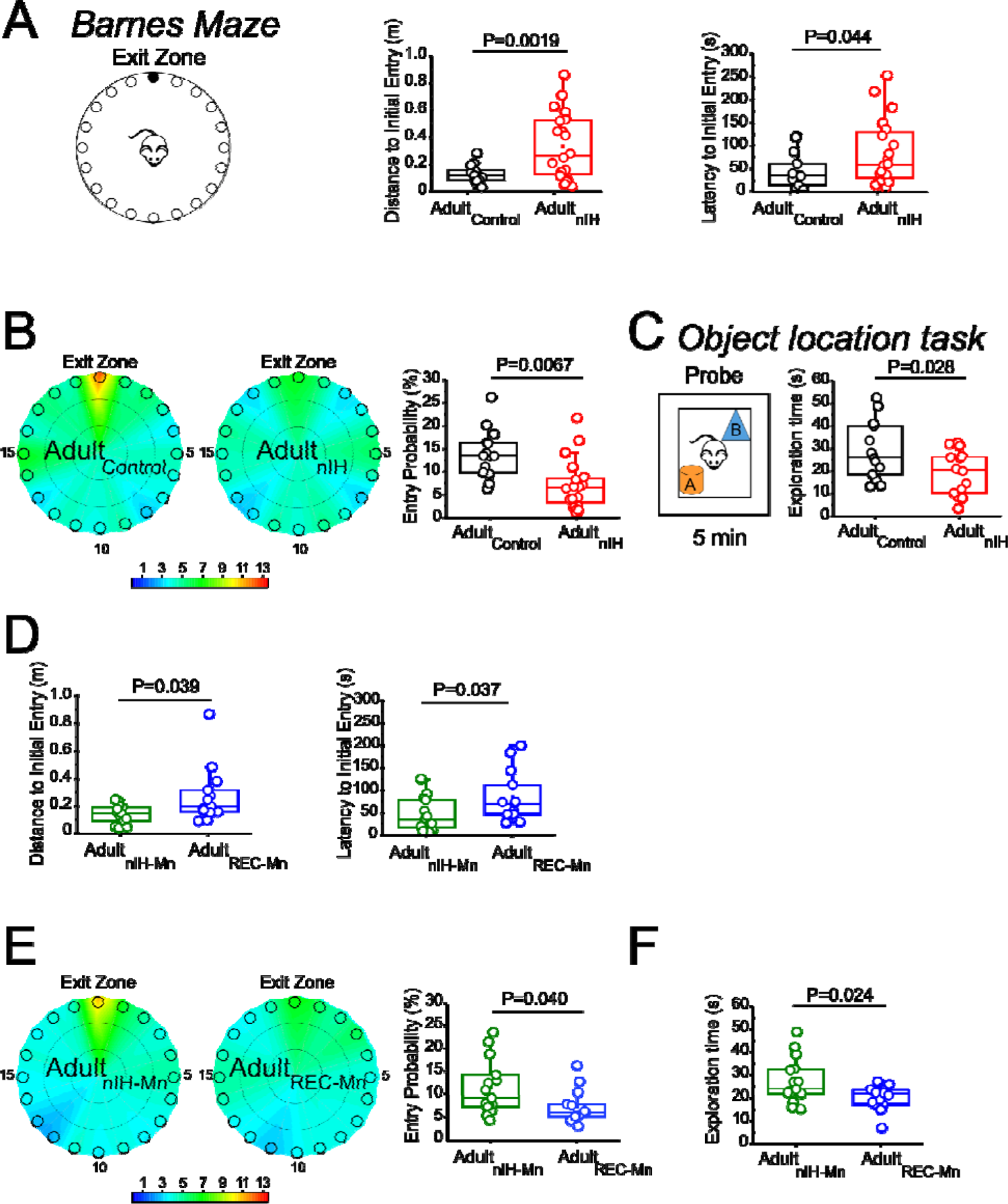
Adult mice exposed to nIH and received MnTMPyP after nIH exhibit spatial memory impairment. **A**. (*left*) Barnes maze diagram. (*middle*) During probe trial, the distance traveled prior to initial entry into the exit zone was greater in IH compared to adult control (t=3.51, df=22.57; P=0.0019). (*right*) The latency finding the exit zone was shorter in Adult control vs Adult_nIH_ (t=2.10, df=30.54; P=0.044). **B**. (*left*) Heat maps of the mean entry probability in control and Adult_nIH_. (*right*) Entry probability into the exit zone is greater in control compared to Adult_nIH_ (t=2.92, df=28.45; P=0.0067). **C**. (*left*) Object location task diagram. (*right*) During the probe trial, control mice exhibit greater exploration time compared to nIH. (t=2.18, df= 24.53;P=0.028). **D**. (*left*) During the probe trial in Barnes maze, the distance to traveled to initial entry into the exit zone was higher in Adult_REC-Mn_ compared to Adult_nIH-Mn_ (t=2.27, df=14.00; P=0.039). (*right*) The latency to finding the entry was smaller in Adult_nIH-Mn_ vs Adult_REC-Mn_ (t=2.23, df=19.37; P=0.037). **E**. (*left*) Heat maps of the mean entry probability in Adult_nIH-Mn_ and Adult_REC-Mn_. (*right*) The entry probability into the exit zone is greater in Adult_nIH-Mn_ compared to Adult_REC-Mn_ (t=2.17, df=23.91; P=0.040). **F**. Adult_REC-Mn_ mice spend more time to the moved object compared to Adult_REC-Mn_ (t=2.52, df=14; P=0.024). The values indicate mean ± S.E. The analysis was performed for B-D, G-J and L using unpaired two-tailed t-test with Welch’s correction.

During the open field session in the object location task, Adult_control_ (N=14) and Adult_nIH_ (N=20) exhibited similar velocities, distance travelled, time in the periphery and time in the center (**Supplement 5**), suggesting no locomotor differences between groups. During familiarization session, control and Adult_nIH_ both exhibited a progressive decrease in exploration times (**Supplement 6**) and similar exploration times with both objects (**Supplement 6**). These data suggested that both groups became learned the location of both objects and did not have preference for either of the objects tested. During the probe trial of the Object location task, control mice exhibit greater exploration time compared to nIH (Figure 5C: control =28.81±3.38s and nIH=19.42±2.63s; P=0.028) We sought to determine whether adult behavioral performance was impacted by MnTMPyP administration during nIH (i.e., Adult_nIH-Mn_) or by MnTMPyP administration following nIH (i.e., Adult_REC-Mn_). Both Adult_nIH-Mn_ (N=16) and Adult_REC-Mn_ (N=13) exhibited progressive reductions in distance traveled across training session without changed velocities (**Supplement 7**). This corresponded with progressive improvement exiting the Barnes maze during the three training sessions (**Supplement 4**). During the probe trial, the initial distance (Figure 5D (*left*); Adult_nIH-Mn_: 0.13 ± 0.016 m versus Adult_REC-Mn_: 0.27± 0.05 m, P=0.039) and latency to initial entry (Figure 5D (*right*); Adult_nIH-Mn_: 43.83± 9.03 s versus Adult_REC-Mn_: 84.56±15.79s, P=0.037) into the exit zone were less in Adult_nIH-Mn_ when compared to Adult_REC-Mn_. Adult_nIH-Mn_ also showed a greater entry probability into the exit zone when compared to Adult_REC-Mn_ (Figure 5E; Adult_nIH-Mn_: 11.45±1.52% versus Adult_REC-Mn_: 7.43 ± 5.18%, P=0.040).

During the open field session of the object location task, Adult_nIH-Mn_ (N=16) and Adult_REC-Mn_ (N=13) exhibited similar velocities, distance travelled, time in the periphery and time in the center (**Supplement 8**), suggesting no locomotor differences between groups. During familiarization, Adult_nIH-Mn_ and Adult_REC-Mn_ both also exhibited a progressive decrease in exploration times (**Supplement 5**) and similar times exploring both objects (Supplemental 5). However, during the probe trial of the Object location task, exploration time for the moved object (i.e., object B) was greater in the Adult_nIH-Mn_ when compared to Adult_REC-Mn_ (Figure 5F; Adult_nIH-Mn_: 27.21 ± 2.51s vs Adult_REC-Mn_: 20.44±1.31, P=0.024)

## Discussion

Intermittent hypoxia can be experienced across a lifetime. While most-prominently associated with untreated sleep apnea in adolescents (Narang & Mathew, 2012) and adults (Ramirez et al., 2013), intermittent hypoxia may also be experienced by children suffering from autonomic dysautonomias, such as that observed in Rett Syndrome and other conditions (Carroll et al., 2015; Glaze, Frost, Zoghbi, & Percy, 1987; Ramirez et al., 2020). In perinatal life, intermittent hypoxia may be experienced embryonically when sleep apnea is left untreated during pregnancy (Dominguez, Street, & Louis, 2018)); whereas, in premature infants, it occurs with apneas of prematurity (Martin, Wang, Koroglu, Di Fiore, & Kc, 2011). Each of these conditions have associations with neurocognitive impairment (Poets, 2020; Ramirez et al., 2013; Slattery et al., 2023), leading to the hypothesis that IH exposure, independent of life stage, impacts neurophysiology and behavior. Over the past three decades, the majority of research examining this hypothesis has been almost exclusively focused on examining how IH impacts neurophysiology in the adult brain. Here, we demonstrate a role for the nIH-dependent pro-oxidant state to remodel NMDAr subunit composition and to cause deficits in synaptic plasticity in the hippocampus. These phenomena are evident immediately follow nIH and persist later in life where they coincide with neurobehavioral deficit. Importantly, our results also show that antioxidant administration has a discrete window to be effective in preventing the nIH-dependent phenomena observed. As our study used subjects with no known genetic predisposition for neurophysiological or behavioral deficits, our findings underscore the critical importance for stable oxygen homeostasis and suggest that neonatal intermittent hypoxia is an independent risk factor for negatively impacting neurophysiological development and behavior later in life.

Recent advancements in understanding how IH impacts synaptic plasticity and neurogenesis in the adult hippocampus has established a mechanistic framework by which IH-dependent HIF1a signaling promotes a pro-oxidant condition (Arias-Cavieres et al., 2020; Chou et al., 2013; Khuu et al., 2021), to suppress neurogenesis (Khuu et al., 2021; Khuu et al., 2019) and impair synaptic plasticity (Arias-Cavieres et al., 2021; Arias-Cavieres et al., 2020; Khuu et al., 2019) as these phenomena are mitigated with antioxidant treatment (Arias-Cavieres et al., 2021; Arias-Cavieres et al., 2020; Khuu et al., 2019). nIH also promotes a pro-oxidant state that coincides with increased nuclear HIF1a signaling and upregulation in NOX isoforms known to be increased by IH-dependent HIF1a activity in the adult (Arias-Cavieres et al., 2020; Chou et al., 2013; Peng et al., 2014; Peng et al., 2006). However, rather than downregulating the obligatory subunit of the NMDAr, as observed in adult hippocampus (Arias-Cavieres et al., 2021; Arias-Cavieres et al., 2020), it appears that the IH-dependent pro-oxidant state perturbs the normal transition of predominant expression of GluN2B subunit to GluN2A in the in the neonatal hippocampus.

The conversion of GluN2 subunit predominance from GluN2B to GluN2A in the brain normally occurs during early postnatal development. In the primary visual cortex of dark-reared animals, GluN2 subunit remodeling is rapidly precipitated by a single one-hour exposure to a visual stimulus (Philpot, Sekhar, Shouval, & Bear, 2001; Quinlan, Philpot, Huganir, & Bear, 1999) demonstrating that GluN2 subunit dominance can be shaped by early life experience. Consistent with this view, experiencing short repeated oscillations in oxygenation in early postnatal effectively influences the transition in NR2 subunit identity within hippocampus. Additionally, our findings indicate that this experience-dependent phenomenon occurring in early life persists to affect hippocampal synaptic properties and associated behavioral outcomes later in life.

As GluN2 subunit composition dictates the biophysical, pharmacological and signaling properties of the NMDAr (Cull-Candy & Leszkiewicz, 2004; Traynelis et al., 2010; Vyklicky et al., 2014), which in turn, influences synaptic timing and summation properties (S. S. Kumar & Huguenard, 2003; Lei & McBain, 2002; Paoletti et al., 2013), the differences in subunit composition would be predicted to impact NMDAr dependent physiology. Indeed, the nIH-dependent deficits to LTP appeared to result from changed GluN2 subunit composition as LTP following nIH was prevented by blocking the GluN2B containing NMDAr and yet, was insensitive to blockade of the GluN2A containing NMDAr. As CAMKII activity is associated with GluN2B containing NMDAr (Hosokawa et al., 2021; Paoletti et al., 2013; Tang et al., 2020), the continued predominance of GluN2B activity may increase CAMKII activity and subsequently affect hippocampal neurophysiology and associated behaviors. However, only MnTMPyP treatment during nIH effectively mitigated the downregulation of GluN2A receptors while also preventing the nIH-dependent effects on LTP and behavioral performance. Thus, while further resolution is required to determine how GluN2B gain of function and GluN2A loss of function each contribute to the emergence and persistence of neurophysiological deficits observed following nIH, our findings with MnTMPyP suggest that GluN2A subunit downregulation is a significant factor contributing to deficits in response to nIH.

Our experiments using MnTMPyP demonstrated that the ability to mitigate the pro-oxidant state during nIH effectively prevents the immediate and long-term consequences of nIH on the hippocampus and behavioral performance. However, MnTMPyP administration following nIH neither mitigated the downregulation of GluN2A nor the impaired synaptic plasticity. When compared to adult mice that received MnTMPyP during nIH, adult mice treated with MnTMPyP following nIH also appeared to have memory deficits as indicated by performance in both the Barnes maze and object location task. Additionally, protein analysis in adult hippocampal tissue following nIH indicated that neither nuclear HIF1a nor NOX isoforms are elevated relative to that in adult mice left in room air. Together these findings strongly suggest that data suggest that while the pro-oxidant state that correlates with increased HIF1a signaling and elevated NOX enzymes may initiate changes in the neonatal hippocampus, the persistent effects of nIH on the hippocampus and behavior later in life are not driven by a persistent elevation in signaling or activity related to these proteins and the pro-oxidant state. Rather, it appears that nIH-mediated pro-oxidant activity initiates a lasting program that remodels hippocampal NMDAr identity and synaptic plasticity to cause behavioral impairments later in life. This lasting program activated by nIH has yet to be determined.

Despite the well-recognized clinical importance of stable oxygenation in early perinatal life, an understanding of the mechanistic role that intermittent hypoxia has on neurodevelopmental outcomes is very limited when compared to the depth of knowledge into the genetic and molecular determinants of intellectual disability. Advancements toward understanding and addressing the consequences of mutations such as SYNGAP1 (Clement et al., 2012; Hamdan et al., 2009; Yang et al., 2023; Zoghbi & Bear, 2012), Fragile X Messenger Ribonucleoprotein 1 (D’Incal, Broos, Torfs, Kooy, & Vanden Berghe, 2022; Leal, Comprido, & Duarte, 2014; Monday, Kharod, Yoon, Singer, & Castillo, 2022), and MECP2 (Banerjee, Miller, Li, Sur, & Kaufmann, 2019) continue to grow. However, as the severity of such neurocognitive deficits often varies among individuals with such genetic predispositions, the extent to which environment interacts with genetic predisposition contribute to augment the severity of neurocognitive deficits is an important issue to be resolved by future work.

## Conflict of Interest

The authors declare no competing financial interests.

## Funding Sources

This work was supported by NIH PO 1 HL 144454 (AJG), NIH R01 NS10742101 (AJG)

---

**Supplement 1:** Control and nIH body mass at postnatal ages used.

| Postnatal Day | Control (N) | nIH and nIH<sub>Saline</sub> (N) | nIH<sub>Mn</sub> (N) | nIH<sub>REC-Mn</sub> (N) | P value |
| --- | --- | --- | --- | --- | --- |
| 4-5 | 2.59±0.06 (42) | 2.48±0.08 (38) | 2.43±0.08 (36) | 2.52±0.10 (13) | P=0.50 |
| 14-15 | 6.17±0.12 (42) | 6.24±0.27 (38) | 5.88±0.14 (36) | 6.98±0.019 (13) | P=0.29 |
| 55-60 | 28.32±0.78 (14) | 29.49±0.66 (20) | 28.78±0.60 (16) | 30.11±0.66 (13) | P=0.43 |

**Supplement 2:**
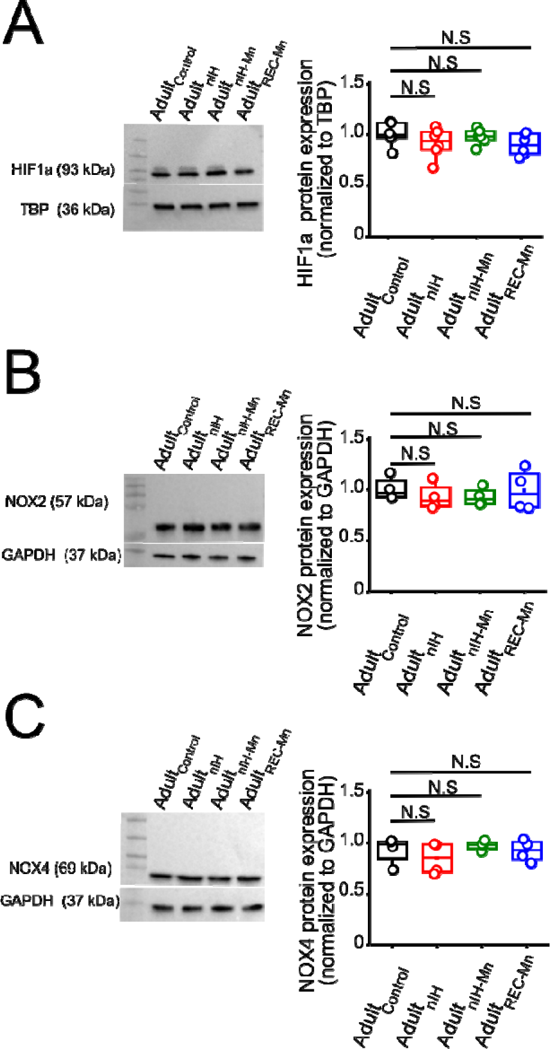
Adult mice exposed to IH do not have increased HIF1a, NOX2 and NOX4 expression. **A**. (*left*) Representative blot of nuclear HIF1a performed from adult mice unexposed (Adult_control_), adult mice were exposed to nIH (Adult_nIH_), adult mice were that received MnTMPyP during nIH exposure (Adult_nIH-Mn_) and adult mice that receive MnTMPyP after nIH exposure (Adult_REC-Mn_). (*right*). Quantification of nuclear HIF1a expression from adult control, Adult_nIH_, Adult_nIH-Mn_ and Adult_REC-Mn_. (one way ANOVA, F_(3,20)_=1.14; P=0.35, N=6). **B**. (*left*) Immunoblot of NOX2. (*right*) No significant differences was found in hippocampal homogenate from adult control, Adult_nIH_, Adult_nIH-Mn_ and Adult_REC-Mn_ (one way ANOVA, F_(3,12)_=10.35; P=0.78, N=4). **C**. (*left*) Representative image of NOX4. (*right*) Comparison of NOX4 expression between adult control, Adult_nIH_, Adult_nIH-Mn_ and Adult_REC-Mn_. (one way ANOVA, F_(3,12)_=0.79; P=0.51,N=4). The box plot parameters indicate mean ± S.E. The analysis was performed for A-C using one-way ANOVA followed by Bonferroni post hoc. N.S= no significant.

**Supplement 3:** Velocity and distance in Adult_control_ and Adult_nIH_/Adult_nIH-Saline_ during Barnes maze training sessions.

|  | **Adult<sub>control</sub>** (N=14) | **P value** | **Adult<sub>nlH</sub> /Adult<sub>nlH-Saline</sub>**<br>(N=20) | **P value** |
| --- | --- | --- | --- | --- |
| Velocity (m/s) | Session 1:0.058 ±0.003<br>Session 2:0.053 ±0.005<br>Session 3:0.049 ±0.005 | P=0.70 | Session 1:0.06±0.006<br>Session 2:0.054±0.005<br>Session 3:0.055±0.005 | P=0.82 |
| Distance traveled (m) | Session 1:18.53 ±2.05<br>Session 2:8.68 ±1.20<br>Session 3: 4.8 ±1.48 | P<0.001 | Session 1:20.58±2.56<br>Session 2:12.27±1.53<br>Session 3:7.36±1.73 | P=0.006 |

**Supplement 4:**
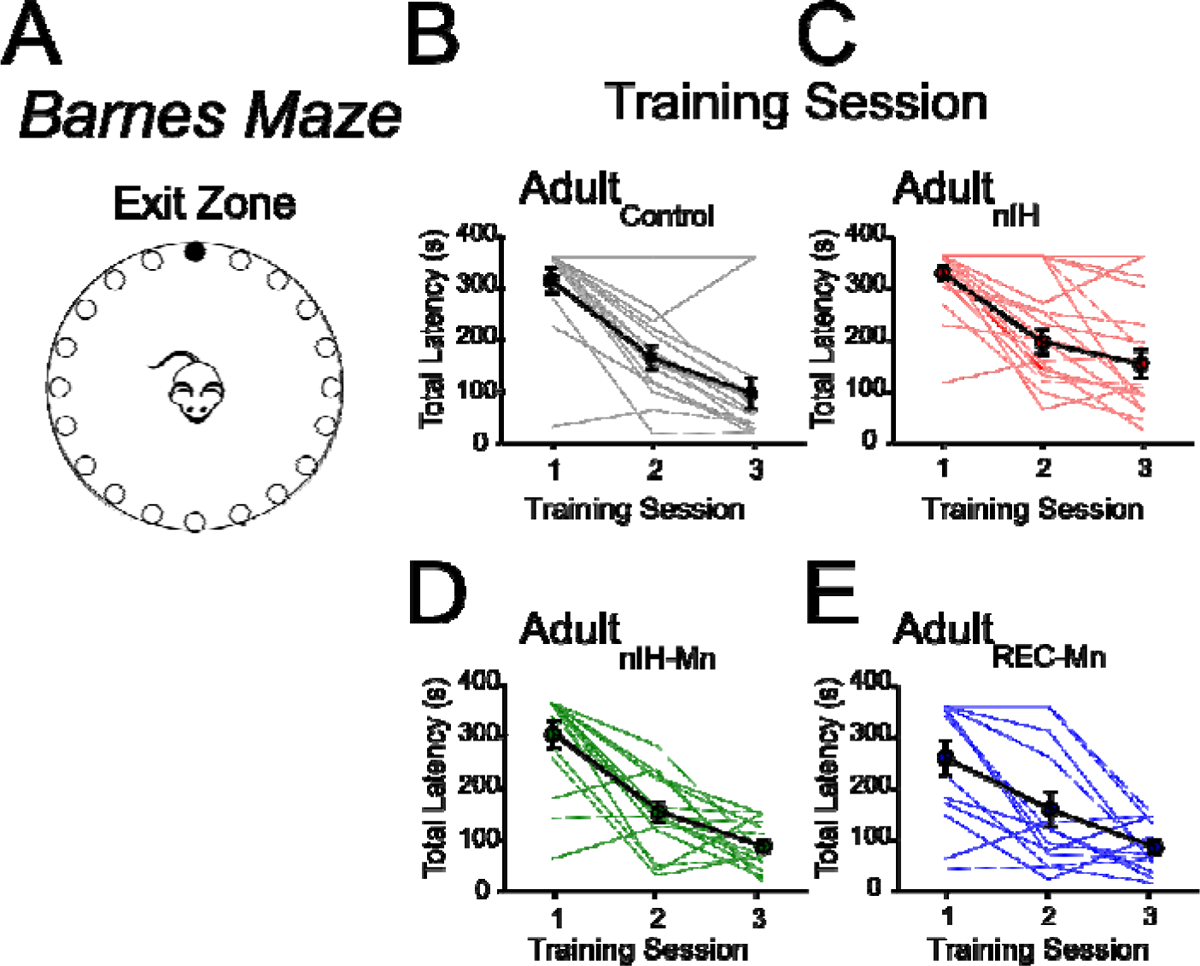
Behavioral performance during training in the Barnes maze is evident across all experimental groups. **A**. Barnes maze diagram. B-**C**. During the training session, the total latency average to the exit hole decreased over three training session in Adult_control_ (F_(2,39)=18.47_, P<0.0001, N=14) and Adult_nIH_ (F_(2,57)=10.98)_, P=0.0015, N=20). Black line represents average latency per trial whereas gray and red lines represent individual performance during training. D-**E**. Black lines represents average latency per trial whereas green and blues lines represent and individual latency during training ((Adult_nIH-Mn_) F_(2,45)=31.55_, P<0.0001, N=16) and Adult_REC-Mn_ (F_(2,36)=9.14_, P=0.0006, N=13). The values indicate mean ± S.E. The analysis was performed for B-E using one-way ANOVA followed by Bonferroni post hoc.

**Supplement 5:** Open field locomotor activity in Adult_control_ and Adult_nIH_/Adult_nIH-Saline_.

|  | **Adult<sub>control</sub>** (N=14) | **Adult<sub>nIH</sub>/Adult<sub>nIH-Saline</sub>** (N=20) | **P value** |
| --- | --- | --- | --- |
| Velocity (m/s) | 0.05 ± 0.003 | 0.05 ±0.003 | P=0.48 |
| Distance traveled (m) | 36.16±2 | 31.13 ±2.08 | P=0.46 |
| Time in the periphery (s) | 361.3 ± 21.26 | 317.5± 23.16 | P=0.17 |
| Time in the center (s) | 237.8±21.22 | 281.6 ±23.13 | P=0.15 |

**Supplement 6:**
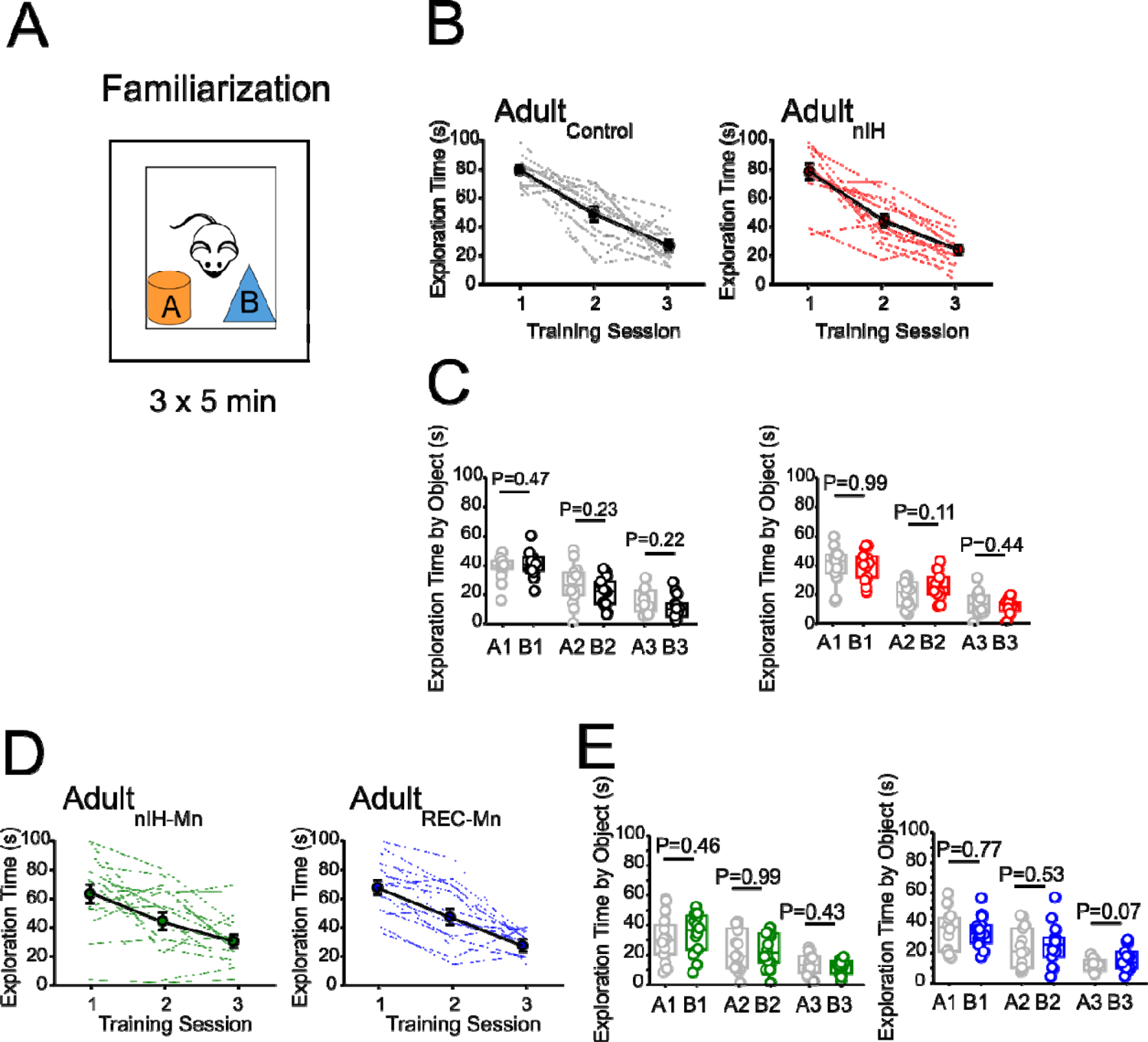
Exploration of objects during Familiarization is similar in all experimental groups. **A**. Object location diagram. **B**. In the familiarization phase, the exploration time in control ((*left*) F_(2,36)=39.05)_, P<0.001; N=14) and IH ((*right*) F_(2,27)=18.33)_, P<0.0001; N=20) decrease across to the three training session. Black line represents average latency per trial whereas grey and red represent individual performance during training. **C**. (*left*) Control mice have similar exploration time with the two objects in the first (t=0.80, df=23.94; P=0.47), second (t=1.52, df=22,63; P=0.23) and third (t =1.36, df=24; P=0.22) session. (*right*) IH mice explored the two-object similar time in the first session (t=0.06, df=17.08; P=0.99), second session (t=1.34, df=17.79; P=0.11) and third session (t=0.06, df=15.92; P=0.44). **D**. Mice received MnTMPyP during nIH exposure (Adult_nIH-Mn_) and mice received MnTMPyP after nIH exposure (Adult_REC-Mn_) decreased the exploration time across the three training session. Adult_nIH-Mn_ ((*left*) F_(2,39)=6.10_, P=0.0049, N=16) and Adult_REC-Mn_ (right, F_(2,33)=14.33_, P<0.0001, N=13). Black line represents average latency per trial whereas green and blue lines represent individual performance during training. **E**. (*left*) Adult_nIH-Mn_ mice explored the two-object similar time in the first (t=0.74, df=29.62; P=0.46), second (t=0.008, df=29.09; P=0.99) and third (t=0.78, df=26.35; P=0.43) sessions. (*right*) Adult_REC-Mn_ mice spent similar time exploring the tow objects in the first (t=0.28, df=26.09; P=0.77), second (t=0.62, df=28.00; P=0.53) and third (t=1.88, df=20.67; P=0.07) sessions. The values indicate mean ± S.E. The analysis was performed for B and D using one-way ANOVA followed by Bonferroni post hoc. The analysis was performed for C and E using Paired-two-sided test.

**Supplement 7:** Velocity and distance of MnTMPyP treated mice during Barnes maze training sessions.

|  | **Adult<sub>nIH-Mn</sub> (N=16)** | **P value** | **Adult<sub>REC-Mn</sub> (N=13)** | **P value** |
| --- | --- | --- | --- | --- |
| Velocity (m/s) | Session 1:0.07 ±0.001<br>Session 2:0.07 ±0.003<br>Session 3:0.064 ±0.008 | P=0.79 | Session 1:0.07±0.02<br>Session 2:0.07±0.004<br>Session 3:0.065±0.012 | P=0.82 |
| Distance traveled (m) | Session 1:18.29 ±1.05<br>Session 2:10.42±1.28<br>Session 3: 5.32 ±0.82 | P<0.001 | Session 1:15.32±2.86<br>Session 2:7.84±1.31<br>Session 3:8.25±2.14 | P=0.043 |

**Supplement 8.** Open field locomotor activity in MnTMPyP treated subjects.

**Supplement 7: Velocity and distance of MnTMPyP treated mice during Barnes maze training sessions.**
|  | **Adult<sub>nlH-Mn</sub> (N=16)** | **Adult<sub>REC-Mn</sub> (N=13)** | **P value** |
| --- | --- | --- | --- |
| Velocity (m/s) | 0.05 ± 0.003 | 0.055 ±0.002 | P=0.33 |
| Distance traveled (m) | 35.21±1.45 | 33.25±1.24 | P=0.35 |
| Time in the periphery (s) | 320.7 ± 26.31 | 304.1± 27.19 | P=0.66 |
| Time in the center (s) | 272±24.65 | 280.2 ±27.12 | P=0.82 |

## Notes

### Competing Interest Statement

The authors have declared no competing interest.

